# Glycation enhances protein association with lipid bilayer membranes

**DOI:** 10.1101/2025.10.30.685514

**Authors:** Beatrice Barletti, Nicoló Paracini, Giovanna Fragneto, Jean-Pierre Alcaraz, Andrew Nelson, Isabelle Vilgrain, Donald K. Martin, Marco Maccarini

## Abstract

Glycation is a non-enzymatic post-translational modification that leads to the formation of advanced glycation end-products (AGEs), which accumulate in the blood-stream under chronic hyperglycemia and are implicated in diabetes-related pathologies. While glycated proteins such as albumin or hemoglobin are widely used as biomarkers for glycemic control, the structural and chemical changes induced by glycation may also alter their interactions with lipid interfaces, including cellular membranes and lipoproteins, potentially affecting their biological distribution and diagnostic detectability.

In this study, we investigated how glycation influences the interaction of bovine serum albumin (BSA) with supported lipid bilayers (SLBs) of different compositions, used as model systems to replicate the diversity of membrane surface charges and fluidity. Using neutron reflectometry (NR), we compared the membrane association of BSA and a chemically-enhanced glycated form of BSA (gBSA), focusing on nanostructural changes at the bilayer interface. Our results showed negligible interaction of either proteins with zwitterionic or cationic membranes. In contrast, both BSA and gBSA exhibited significant binding to negatively charged bilayers, with glycation significantly amplifying this interaction. Quantitatively, the membrane-associated protein volume fraction increased from 0.11 (BSA) to 0.17 (gBSA), suggesting that glycation modifies the protein’s surface properties in ways that promote stronger lipid interactions with negatively charged membranes.

These findings suggest that glycation not only affects protein structure but also modulates protein–membrane affinity in a lipid-dependent manner. This has important implications for the bioavailability and behavior of glycated albumin in the bloodstream, potentially influencing the accuracy of clinical assays and contributing to membrane-related pathophysiology in diabetes. Our work highlights the need for a deeper understanding of glycation-induced changes in protein-lipid interactions and their consequences for biomarker reliability and disease mechanisms.

**Graphical TOC Entry:** 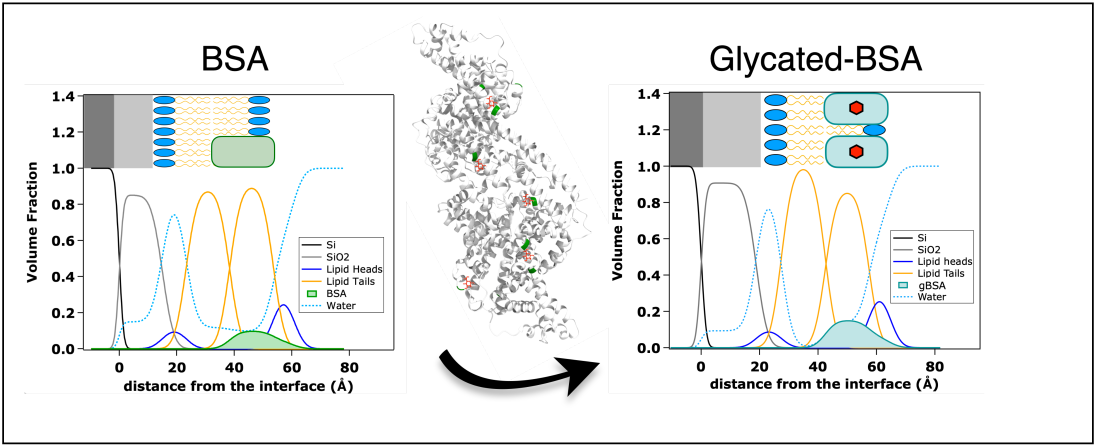

## Introduction

Most plasma proteins are subject to glycation, a non-enzymatic reaction in which reducing sugars, such as glucose, covalently bind to free amino groups, typically at the N-terminus or at the side chains of lysine and histidine residues.^1–5^ This process, also known as the Maillard reaction,^6^ occurs in the bloodstream and differs from glycosylation, which is an enzyme-catalysed modification that primarily takes place in the endoplasmic reticulum and Golgi apparatus. ^7^ Glycation results in the formation of advanced glycation end-products (AGEs),^8^ which accumulate over time and are implicated in several pathological conditions, including type II diabetes mellitus,^1,5,9–11^ cancer,^12–14^ and age-related diseases. ^15–17^

Glycated proteins and AGEs provide an integrated snapshot of blood glucose exposure over defined time frames, making them valuable biomarkers not only for monitoring glycemic control^18^ but also for predicting the risk of diabetes-related complications through their quantification in blood or tissues. ^19^ Plasma and extracellular matrix proteins are particularly prone to glycation due to their continuous exposure to circulating glucose. Among these, hemoglobin A1c (HbA1c) is the most commonly used marker, reflecting average glucose levels over a long period of 8–12 weeks.^20,21^ In contrast, glycated human serum albumin (HSA) reflects shorter-term glycemic changes over a period of 1–3 weeks.^17,22^ Unlike HbA1c, glycated albumin can be measured without the need for fasting, offering a practical advantage for clinical screening. Moreover, it has also been shown to be more effective than HbA1c in detecting prediabetes in non-obese individuals, where traditional markers may fail.^23^ Therefore, the combined use of HbA1c and glycated albumin has been proposed to improve diagnostic sensitivity, particularly in early or borderline cases.^19,23^

Glycated albumin levels are considered unaffected by recent food intake, allowing for non-fasting blood collection, which is a feature that enhances its practicality as a clinical biomarker. However, although glycated albumin is primarily studied in the context of glycemic monitoring, its structural modifications may have broader biological implications. Because glycosylated proteins have been shown to be associated with lipid particles in the bloodstream such as HDL and LDL,^24–29^ it is reasonable to expect that protein glycation might also play a role in the interaction with lipoproteins. However, the literature in this field remains limited. Indeed, no targeted studies have explicitly tested the extent to which glycated albumins bind to, or are altered by, the presence of lipoproteins. This gap in knowledge raises the possibility that glycation may modulate how plasma proteins interact with lipid environments, in the bloodstream, on lipoprotein particles, or at cell membranes, depending on the extent and distribution of glycation sites.

To explore this hypothesis, in this study we investigated the nanostructural effects of bovine serum albumin (BSA) and a chemically enhanced glycated form of BSA on artificial lipid membranes. Since serum-derived BSA may exhibit a baseline level of glycation, we refer to the modified form as enhanced glycated BSA (gBSA) throughout this study. BSA was chosen due to its structural similarity to HSA, its extensive use as a model protein in glycation research and the availability of well-established glycation protocols. By comparing BSA and gBSA, we aimed to highlight the influence of glycation on protein–lipid interactions. Supported lipid bilayers (SLBs) composed of four distinct lipid mixtures were used to replicate different membrane environments, enabling controlled modulation of surface charge and fluidity. The structural changes in the membrane upon protein interaction were characterized using neutron reflectometry (NR), a powerful technique capable of resolving the structure of planar stratified systems with subnanometer precision. This approach enabled us to investigate how enhanced glycation alters protein-lipid interactions and how specific biophysical membrane properties influence these interactions.

Importantly, if glycated proteins such as gBSA exhibit affinity for lipid membranes or lipoprotein particles, this behavior could impact their bioavailability in the bloodstream and the sensitivity or accuracy of diagnostic assays based on their serum concentration. Understanding these interactions is therefore not only mechanistically relevant but also critical for evaluating the robustness of glycated albumin as a clinical biomarker in diabetes monitoring. Additionally, exploring how glycated proteins interact with cell membranes could open new avenues for understanding the molecular mechanisms driving glycation-related diseases. Furthermore, this line of research may reveal new therapeutic approaches aimed at preventing or reversing the deleterious effects of glycation on cellular membranes, potentially offering new strategies to mitigate the complications of diabetes and other diseases associated with elevated AGEs.

## Materials and methods

### Materials

Bovine Serum Albumin (BSA) was purchased from Sigma-Aldrich (Darmstadt, Germany) in the form of lyophilized powder product number A8806-1G (CAS 9048-46-8); D-(+)-Glucose was purchased from Sigma-Aldrich (Darmstadt, Germany) (CAS 50-99-7); Tris buffered saline pH 7.6 buffer was prepared using the BioUltra tablets from Sigma-Aldrich (Darmstadt, Germany). 1-palmitoyl-2-oleoyl-glycero-3-phosphocholine (*C*_42_*H*_82_*NO*_8_*P* , 16:0-18:1 PC, POPC, 760.076 g/mol, CAS: 26853-31-6), 1-palmitoyl-2-oleoyl-sn-glycero-3-phospho-L-serine sodium salt (*C*_40_*H*_75_*NO*_10_*PNa*, 16:0-18:1 PS, POPS, 783.988 g/mol, CAS: 321863-21-2), 1,2-dioleoyl-3-trimethylammonium-propane chloride salt (*C*_42_*H*_80_*NO*_4_*Cl*, 18:1 TAP, DOTAP, 698.542 g/mol, CAS: 132172-61-3), brain porcine sphingomyelin (*C*_41_*H*_83_*N*_2_*O*_6_*P* , SM, 760.223 g/mol, CAS: 383907-91-3) and ovine wool cholesterol (*C*_27_*H*_46_*O*, CHOL, 386.654 g/mol, CAS: 57-88-5) for SLBs preparation were purchased from Avanti Polar Lipids (Alabaster, AL, USA). Tris buffer saline (TBS) for protein preparation and formation of SLB was previously degassed in a bath sonicator and filtered with a 0.2 µm filter prior to use in the experiments.

### Glycation of Bovine serum albumin

The glycation of BSA was performed following a protocol employed in previous studies with minor modifications.^30,31^ Glucose was used as the reducing sugar in the reaction, as it is the predominant sugar in blood. To accelerate the reaction, stock solutions of 300 *µ*M BSA and 2 M glucose in 10 *µ*M PBS buffer pH 7.6 were prepared. The stock solutions were filtered to 0.2 *µ*m. To perform the glycation, 500 *µ*L of BSA and 500 *µ*L of glucose stock solutions were mixed obtaining final concentrations of 150 *µ*M and 1 M for BSA and glucose respectively. The mixed solution was incubated in the dark for 14 days at 37°C to obtain gBSA. A solution of BSA at concentration 150 *µ*M without glucose was also incubated for 14 days at 37°C to obtain a control reference. Notably, the half-life of BSA under similar in vitro conditions has been reported to be approximately 20 days,^32^ which supports the stability of the protein during the incubation period.

### Supported lipid bilayers preparation

SLBs were prepared using the vesicles fusion technique with lipid vesicles of varying compositions. The vesicles were derived from lipid films consisting of pure, binary, and ternary lipid mixtures, specifically POPC, POPC/POPS (8:2), POPC/DOTAP (7:3), and POPC/SM/CHOL (6:3:1). These films were generated by mixing stock chloroform solutions (1 mg/mL) of each lipid component in the specified molar ratios. The solvent was removed under a gentle nitrogen stream, followed by overnight drying under vacuum to ensure complete evaporation.

The resulting mixed lipid films were then resuspended in milliQ H_2_O achieving a final lipid concentration of 0.2 mg/mL. To facilitate vesicle formation, the suspensions underwent bath sonication for 20 minutes, followed by tip sonication on ice for 15 minutes at 50% amplitude, with 5-second on/off pulses, resulting in vesicles of approximately 100 nm in diameter. SLB formation for NR experiments was achieved via spontaneous vesicles fusion, immediately before injecting the vesicles the sonicated lipid suspension was mixed 1:1 with a 4 mM solution of Ca*Cl*_2_ to yield a final concentration of 0.1 mg/mL lipids and 2 mM Ca*Cl*_2_.

### MALDI-TOF

Matrix-assisted laser desorption/ionization time-of-flight (MALDI-TOF) mass spectrometry was used to characterize changes in the molecular weight (MW) of BSA following glycation. Native and glycated BSA samples were prepared at a concentration of 150 *µ*M and diluted 1:10 in 0.1% trifluoroacetic acid (TFA). Samples were mixed with a matrix solution consisting of *α*-cyano-4-hydroxycinnamic acid (CHCA) and 2,5-dihydroxybenzoic acid (DHB) in equal volumes and spotted onto a MALDI target plate.

Mass spectra were acquired using a MALDI-TOF instrument (Bruker) at the Institute de Biologie Structurale (IBS), Grenoble. Laser power was set to 25% or 40% depending on sample conditions. Data acquisition was performed in linear mode over a mass range of 20–120 kDa. Calibration was carried out using Protein Calibration Standard II (Bruker) under identical conditions. Molecular weight shifts between native and glycated BSA were analyzed to assess glycation-induced modifications.

### Neutron reflectometry (NR)

Neutron reflectometry (NR) measurements were conducted on the FIGARO reflectometer at the Institut Laue-Langevin (ILL), Grenoble, France. ^33^ The instrument operated in time-of-flight mode with a vertical scattering geometry and a horizontal sample orientation. Data were acquired at incident angles of 0.7° and 3.0°, across a wavelength range of 2–20 Å and a Δ*λ/λ* resolution of 7%.

Samples consisted of SLBs assembled on polished silicon blocks (8 × 5 cm^2^, 15 mm thick), with a surface roughness below 5 Å. The bilayers were housed in solid–liquid cells designed for in situ NR measurements. Initial reflectivity profiles were obtained for pristine SLBs. Protein aliquots (1.5 *µ*M) were subsequently introduced into the aqueous subphase and incubated for 5–7 hours before being rinsed with buffer.

To enhance structural sensitivity, contrast variation was applied using isotopically distinct solvent mixtures: 100% D_2_O (SLD = 6.36 × 10*^−^*^6^ Å^-2^), 66% D_2_O/34% H_2_O (“4-matched water”, SLD = 4.00 x 10*^−^*^6^ Å^-2^), 38% D_2_O/62% H_2_O (“silicon-matched water”, SLD = 2.07 x 10*^−^*^6^ Å^-2^), and 100% H_2_O (SLD = –0.56 x 10*^−^*^6^ Å^-2^). Three neutron contrast conditions were recorded for the pristine bilayers, and four were acquired after exposure to the protein, enabling a comprehensive structural analysis before and after interaction. For samples in 100% H_2_O measurements were repeated before and after rinsing to confirm system stability and ensure comparability across contrasts. All contrasts were co-refined to increase the robustness of the resulting structural models.

NR is sensitive to variations in the scattering length density (SLD) profile normal to the interface. The measured quantity is the reflectivity, defined as the ratio of reflected to incident neutron intensities, plotted as a function of the momentum transfer perpendicular to the surface, *Q_z_*, calculated as:

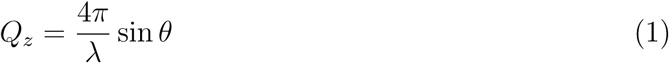

where *λ* is the neutron wavelength and *θ* the incident angle. The SLD, a central parameter in the model, is given by:

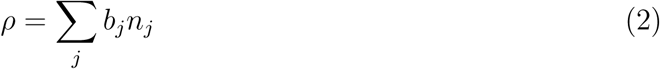

where *b_j_*is the coherent scattering length and *n_j_* the number density of nucleus *j*. Reflectometry data were interpreted through model-based fitting, requiring the construction of a representative interfacial structure composed of discrete, laterally homogeneous slabs. Each slab was described by its SLD, thickness, interfacial roughness, and solvent volume fraction. These parameters define a theoretical reflectivity profile, which was fitted to the experimental data using the RefNX software package.^34^ The reflectivity calculations were based on the optical matrix formalism of Abeles, ^35^ with interface roughness described by the Nevot-Croce approach.^36^ Parameter estimation and uncertainty quantification were performed using a Bayesian framework via Markov Chain Monte Carlo (MCMC) sampling, providing posterior distributions and visual diagnostics through corner plots.

Different structural models were tested to describe the interfacial configuration of the system before and after protein interaction. For the pristine SLB, a five-slab model was employed: SiO_2_, inner headgroup, inner tail, outer tail, and outer headgroup layers, enclosed by semi-infinite silicon and bulk water layers (see Figure S1A in the supporting information (SI)). To simulate protein adsorption, a sixth slab representing a protein layer was added on top of the bilayer (Figure S1B).

In cases where protein insertion into the bilayer was hypothesized, one or more lipid layers were modeled as mixed protein–lipid slabs (Figure S1C–D). Their non-solvent SLDs were calculated as linear combinations of the SLDs of the constituent components, weighted by their volume fractions. Volume fractions for the protein were constrained to remain consistent across contiguous protein-containing slabs to ensure physical continuity.

The SLD of the protein was calculated from its atomic composition and estimated molecular volume using the Biomolecular Scattering Length Density Calculator.^37^ Corrections were applied to account for D/H exchange of labile hydrogen atoms in D_2_O-containing buffers.

Each lipid bilayer dataset was independently fitted using various structural models (schematized in Figure S2 in the SI), with initial model selection guided by the minimization of the normalized *χ*^2^ value. However, the final choice of the most appropriate model was not based solely on the *χ*^2^ criterion. It was complemented by a Bayesian analysis implemented in the RefNX package, which provides posterior distributions of the fitting parameters and insights into their covariance.

In some cases, models with an additional layer yielded slightly lower normalized *χ*^2^ values. However, when evaluating competing models, we also considered the posterior distributions of key structural parameters obtained via Bayesian analysis. If the additional layer’s parameters, such as thickness and hydration, did not exhibit physically meaningful values, the model was considered over-parameterized and less reliable. In such cases, we favored more parsimonious models that avoided overfitting and retained only statistically and physically significant features, better reflecting the underlying physical system.

## Results and Discussion

A MALDI-TOF spectrometer was used to characterize changes in the MW of BSA resulting from the glycation reaction. The spectra of gBSA and native (unmodified) BSA used as control are shown in Figure 1. MALDI-TOF analysis revealed a mass increase in BSA following glycation, with the MW of gBSA measured at 67,224 Da, compared to 66,400 Da for unmodified BSA. Taking into account the MW of glucose (180 Da) and the concurrent loss of water during Schiff base formation (18 Da per glucose molecule), this shift of 824 Da corresponds to the covalent attachment of approximately five glucose molecules. These results indicate that glycation was only partial, as not all available reactive sites were modified. Based on structural analysis, up to 52 lysine and arginine residues in BSA are solvent-exposed and theoretically available for glycation. ^38^ Under highly aggressive glycation conditions, a previous study reported the attachment of up to 48 glucose moieties, suggesting near-complete modification of these available sites.^38^ Additional studies have identified albumin residues that are potentially susceptible to glycation under favorable conditions, ^17^ specifically Lys-524 has been identified as a primary glycation site in BSA. ^39^ In addition, other lysine residues, such as Lys-275, Lys-232, and Lys-396, have also been reported as particularly susceptible to glucose modification.^39^ Despite the moderate glycation yield, in our study the resulting gBSA was sufficient to induce measurable nanostructural effects on SLBs.

**Figure 1:**
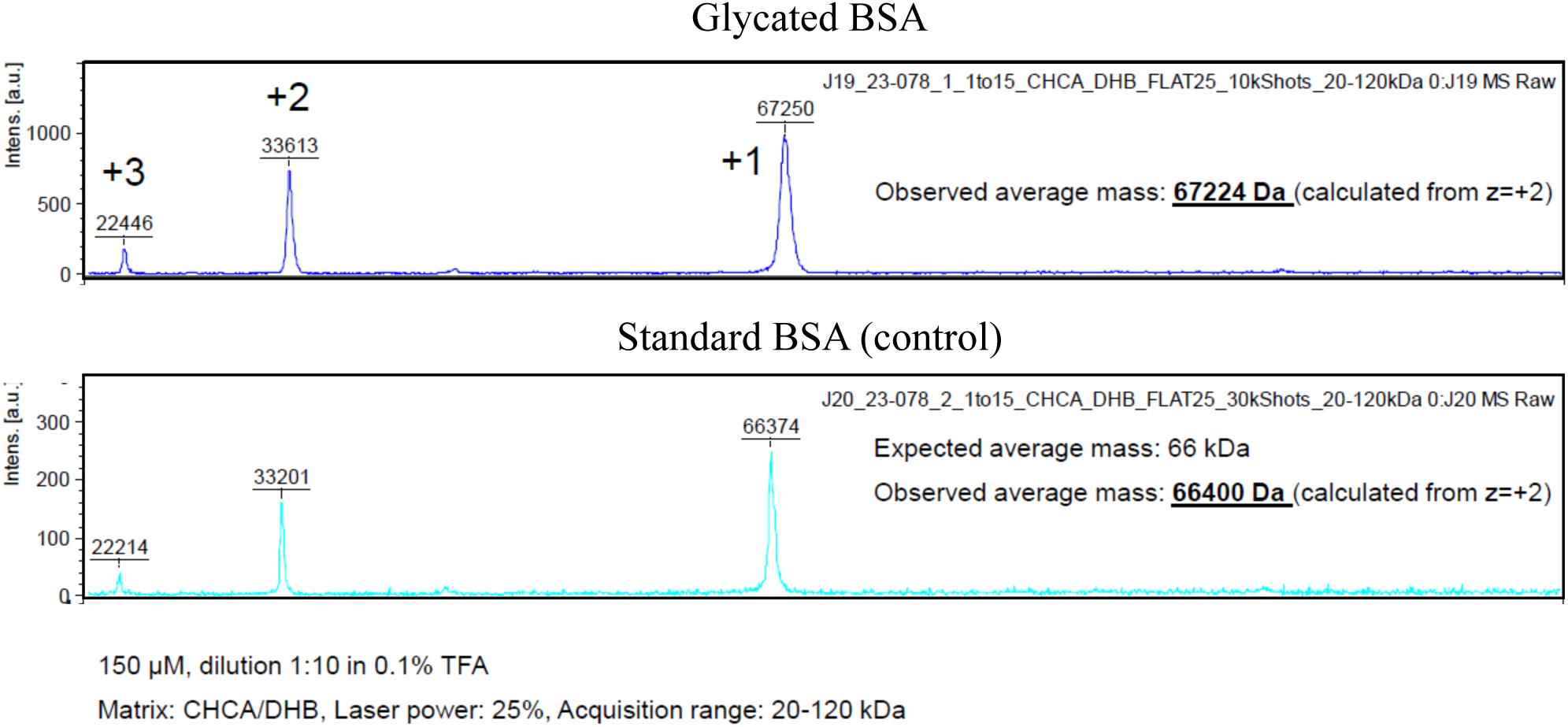
Mass spectra of glycated BSA (gBSA) and standard BSA (control) obtained with MALDI-TOF spectrometer.

Neutron reflectometry was employed to investigate the impact of glycation on protein–lipid interactions. Experiments were performed on all four SLBs adsorbed onto SiO_2_ substrates, both before and after a 5–7 hour exposure to BSA and gBSA. Several structural models were tested against the experimental reflectivity data. The models considered different average spatial arrangements of the protein relative to the bilayer, including: (i) absence of protein near the bilayer, (ii) adsorption of protein onto the bilayer surface without penetration, and (iii) protein adsorption with partial penetration into various regions of the bilayer—ranging from the distal headgroup to the tail region or deeper. For detailed schematics, see Figures S1 and S2 in the SI. In the following, only the structural parameters derived from the best-fitting model are reported.

Figure 2 summarizes the reflectivity profiles in D_2_O and H_2_O for the various SLBs before and after exposure to the two proteins. For SLBs composed of zwitterionic and cationic lipid mixtures, only minor changes in the reflectivity profiles were observed following protein exposure. In contrast, SLBs incorporating anionic lipid compositions exhibited pronounced alterations, indicating a more substantial interaction with the proteins.

**Figure 2:**
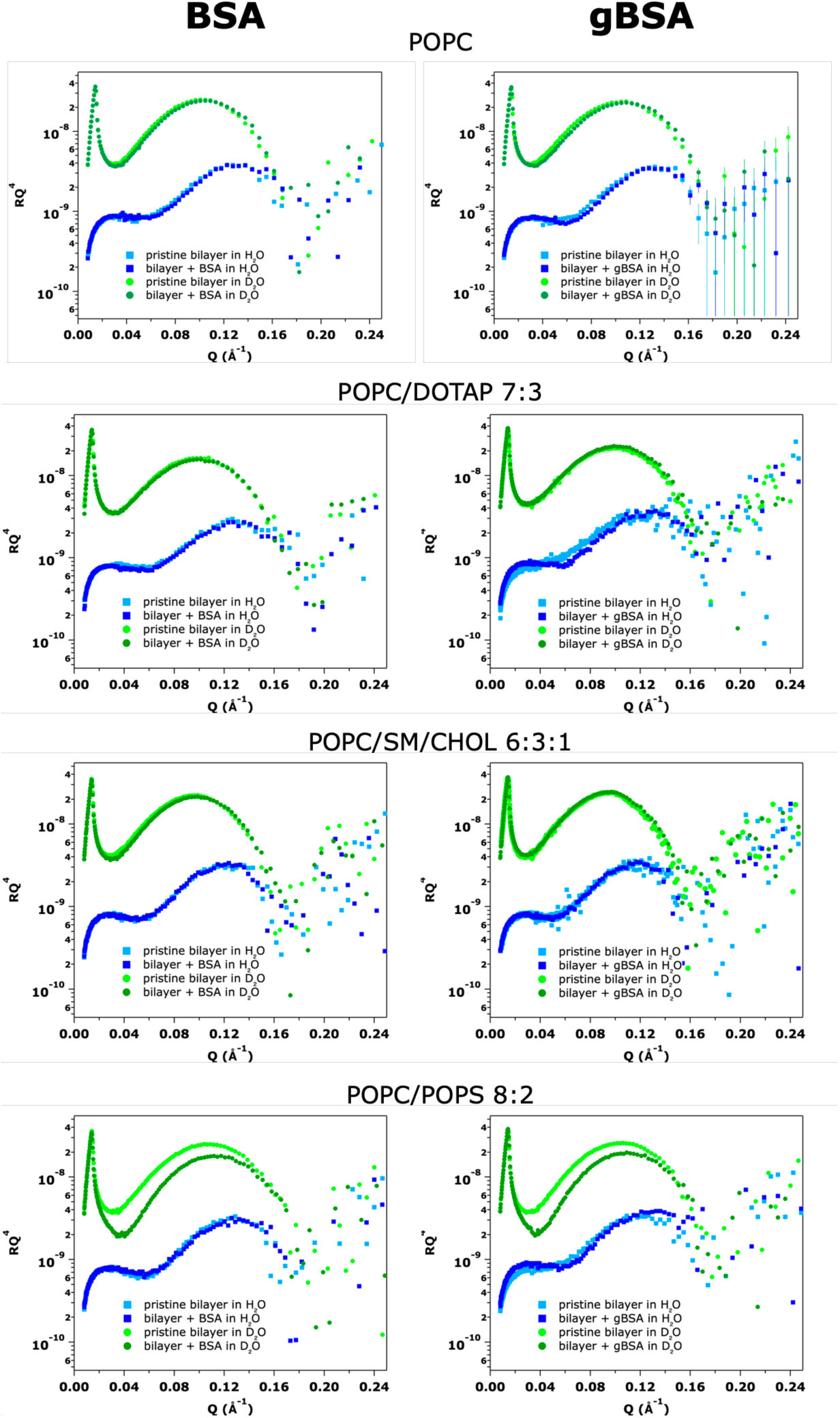
Reflectivity curves in H_2_O and D_2_O contrasts for the four types of SLB before and after the exposure to BSA and gBSA.

Turning to the nanostructural analysis based on model testing, a clear and unambiguous model selection was not achievable for the supported pure POPC bilayer in interaction with either BSA or gBSA. The differences in the reflectivity profiles are too small to allow for a precise identification of the structural changes. Among the tested models, those assuming no interaction and those considering protein penetration into the outer headgroup region, protruding above the bilayer, exhibited the lowest (and similar) normalized *χ*^2^ values (see Table S1 in the SI). The difference in *χ*^2^ between the interaction and non-interaction models was slightly more pronounced for gBSA than for BSA, which may suggest that the interaction model is a better representation of the gBSA data. However, due to the relatively small magnitude of these differences, we cannot conclusively favor the interaction model over the non-interaction model.

As an illustrative examples, Figure S3 in the SI displays the reflectivity profiles and fits at all contrasts, together with the corresponding SLD and volume fraction profiles, for the SLB composed of POPC before and after the exposure to gBSA. The data are modeled assuming protein penetration into the outer headgroup region of the lipid bilayer, accompanied by an additional protruding protein layer above the bilayer. The structural parameters obtained from these fits are reported in table S2 in the SI. While for BSA, we report the best-fitting model describing no interaction with the SLB composed of POPC (see Figure S4 in the SI). The structural parameters obtained from these fits are reported in table S3 in the SI.

The interaction of BSA and gBSA with SLBs composed of POPC/DOTAP (7:3) and POPC/SM/CHOL (6:3:1) had no significant effect on membrane structure (Fig. 2). The fitting analysis revealed that the structural parameters of these SLBs remained largely unchanged before and after exposure to the two proteins. The reflectivity profiles of the POPC/DOTAP (7:3) and POPC/SM/CHOL (6:3:1) SLBs, both before and after BSA and gBSA exposure, along with the best-fit model curves and the corresponding SLD and volume fraction profiles, are provided in the SI (Fig. S5 and S6 for BSA, while Fig. S7 and S8 for gBSA).

For these lipid compositions the model without the presence of the protein was chosen as the model that best fit the data. The structural parameters of the SLBs before and after the exposure to BSA and gBSA are reported in table S3 and table S2 respectively in the SI. With the SLB composed of POPC/POPS 8:2, containing negatively charged lipids, BSA exhibited a significant interaction as it can be seen in Figure 2 bottom panel. This is particularly evident under the D_2_O contrast conditions where marked differences are visible in a large range of Q.

Figure 3 presents the reflectivity profiles of the SLB before and after exposure to BSA, along with the corresponding best-fitting model, scattering length density (SLD), and volume fraction profiles. The best fit corresponds to the model in which BSA partially penetrates the outer leaflet of the bilayer, without the formation of an additional protein layer adsorbed on the membrane surface (see Fig. S2, model 3b). The structural parameters obtained from this fit are summarized in Table 1. Notably, the incorporation of BSA into the outer leaflet is accompanied by increased hydration and interfacial roughness. The fitted thickness of the outer leaflet is 22 Å, slightly smaller than the estimated radius of gyration of BSA (∼ 27 Å).^40^ However, considering the bilayer’s interfacial roughness of approximately 3 Å on each side, the observed thickness of the diffuse layer remains compatible with the protein embedding within the leaflet structure.

**Figure 3:**
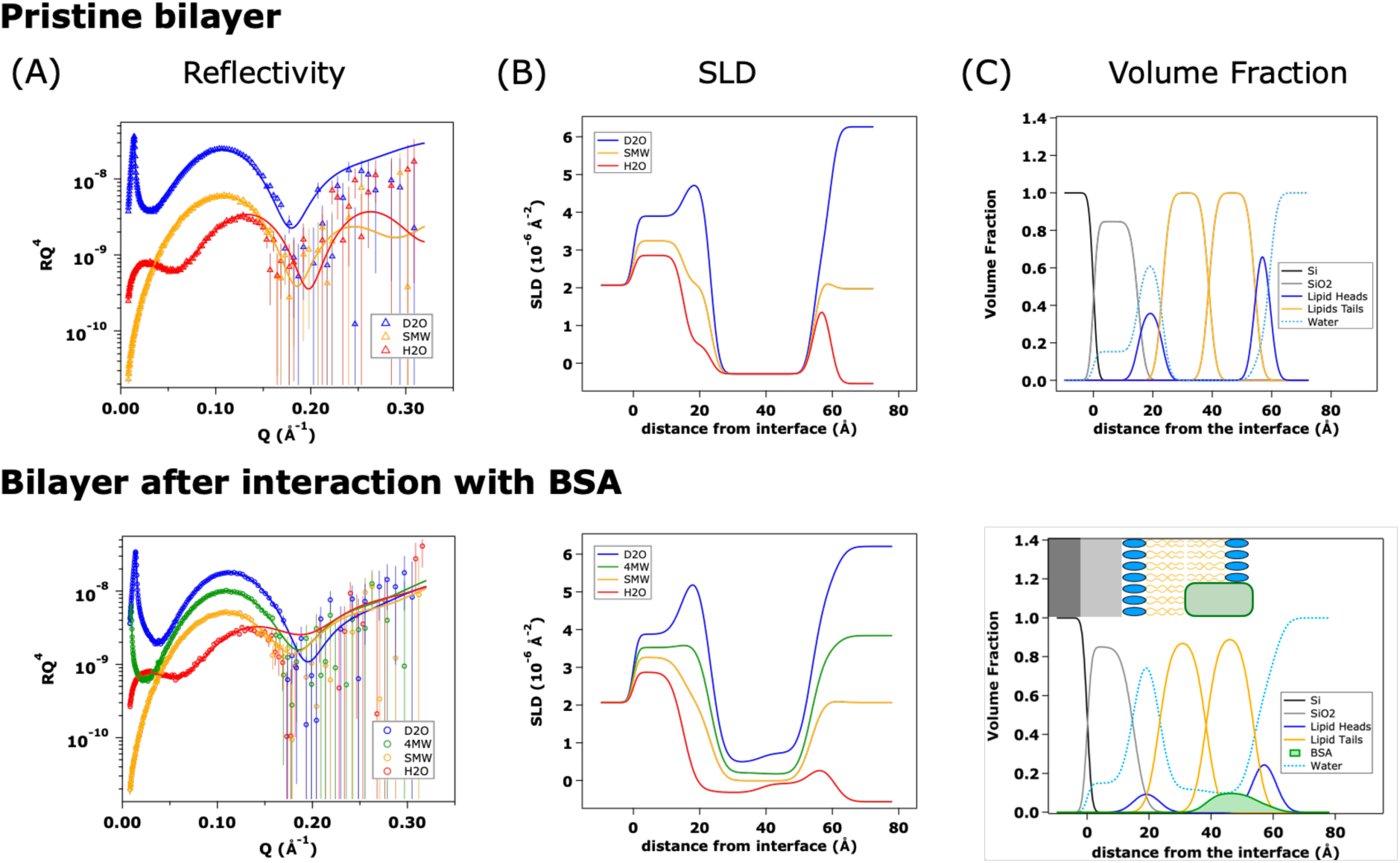
Reflectivity curves and fits at different contrasts (A), the SLDs obtained from the analysis (B) and the corresponding volume fraction distribution (C) of the SLB POPC/POPS 8:2 before and after the exposure to the BSA. Symbols represent the raw data; continuous lines represent the fit. The model that describes best the experimental behaviour is the one representing the penetration of the protein in the outer leaflet of the lipid bilayer with no adsorbed layer of proteins onto the SLB.

**Table 1:** Structural parameters of the SLB POPC/POPS 8:2 before and after the exposure to the BSA. The corresponding fits are in Figures 3. The values without error were kept fixed at their nominal value.

| Layer | Thickness [$\text{\AA}$] | SLD [$\times 10^{-6} \text{\AA}^{-2}$] | Solvent [v/v] | Roughness [$\text{\AA}$] |
| --- | --- | --- | --- | --- |
| **POPC/POPS 8:2** |  |  |  |  |
| SiO<sub>2</sub> | $15 \pm 1$ | 3.47 | $0.15 \pm 0.01$ | $1 \pm 1$ |
| Inner headgroup | $8 \pm 1$ | 2.285 | $0.629 \pm 0.01$ | $2 \pm 1$ |
| Inner chain region | $15 \pm 1$ | -0.282 | $0.00 \pm 0.01$ | $2 \pm 1$ |
| Outer chain region | $15 \pm 1$ | -0.282 | $0.00 \pm 0.01$ | $2 \pm 1$ |
| Outer headgroup | $5 \pm 1$ | 2.285 | $0.17 \pm 0.01$ | $2 \pm 1$ |
| *After exposure to BSA* |  |  |  |  |
| Inner headgroup | $8 \pm 1$ | 2.285 | $0.88 \pm 0.01$ | $3 \pm 1$ |
| Inner chain region | $15 \pm 1$ | -0.282 | $0.12 \pm 0.02$ | $3 \pm 1$ |
| Outer chain region (*) | $15 \pm 1$ | -0.282 | $0.10 \pm 0.02$ | $3 \pm 1$ |
| Outer headgroup (*) | $7 \pm 1$ | 2.285 | $0.63 \pm 0.4$ | $3 \pm 1$ |
| Volume fraction protein | $0.11 \pm 0.04$ | in the slabs labelled with * | | |

Although a similar model incorporating an additional protein layer adsorbed on the membrane surface yielded comparable normalized *χ*^2^ values, we employed a Bayesian posterior analysis to further evaluate model plausibility. Specifically, we examined the posterior distributions of the thickness and solvent volume fraction of the extra layer, which represents the portion of protein protruding above the outer leaflet into the bulk solvent. As shown in Figure S9 (SI), the thickness distribution is sharply peaked near zero, and the solvent volume fraction remains close to one. These features indicate that the additional layer lacks physical meaning and would likely lead to overfitting rather than providing a meaningful improvement to the model. Consequently, the model describing protein penetration into the outer leaflet, without invoking an extra protein layer (see Fig. S1D), was selected as the most appropriate representation of the data. Based on this model, BSA penetrates the bilayer with a volume fraction of 0.11±0.04, accompanied by a moderate increase in leaflet roughness and hydration, potentially indicating partial lipid displacement or rearrangement. Interestingly, despite its net negative charge at physiological pH, BSA showed higher affinity for negatively charged membranes. This counterintuitive behavior aligns with previous studies, suggesting that BSA contains localized cationic patches enabling electrostatic interactions with anionic lipids.^41–43^ Indeed, previous experiments have shown that BSA adsorbs to negatively charged liposomes (e.g. DPPC/DPPG) with adsorption increasing alongside DPPG content, pointing to the role of hydrophobic interactions and maybe positive patches in overcoming electrostatic repulsion.^44^

The gBSA also exhibited a significant interaction with the SLB composed of POPC/POPS 8:2. As for BSA, the best fit was obtained using the model describing the penetration of the protein in the outer leaflet with no adsorbed layer of proteins onto the SLB (see Fig. S2, model 3b). Figure 4 illustrates the reflectivity profiles and fitting curves of POPC/POPS 8:2 SLB, both prior to and following exposure to gBSA, together with the corresponding SLD profiles and the volume fraction distribution of the components. The fitting parameters are presented in Table 2. The resulting model indicates that the protein penetrates the outer leaflet of SLB with a volume fraction of 0.17±0.01. This penetration is accompanied by an increase in the hydration of both the headgroup and tail regions of the lipids, suggesting partial lipid removal, an effect also reported for HSA on PC/PS (8:2) bilayers. ^45^ The increase in hydration is particularly pronounced in the distal (outer) leaflet, likely due to a higher concentration of proteins in this region. This may be attributed to the presence of more hydrophilic moieties on the glycated protein, enhancing its affinity for the membrane interface.

**Figure 4:**
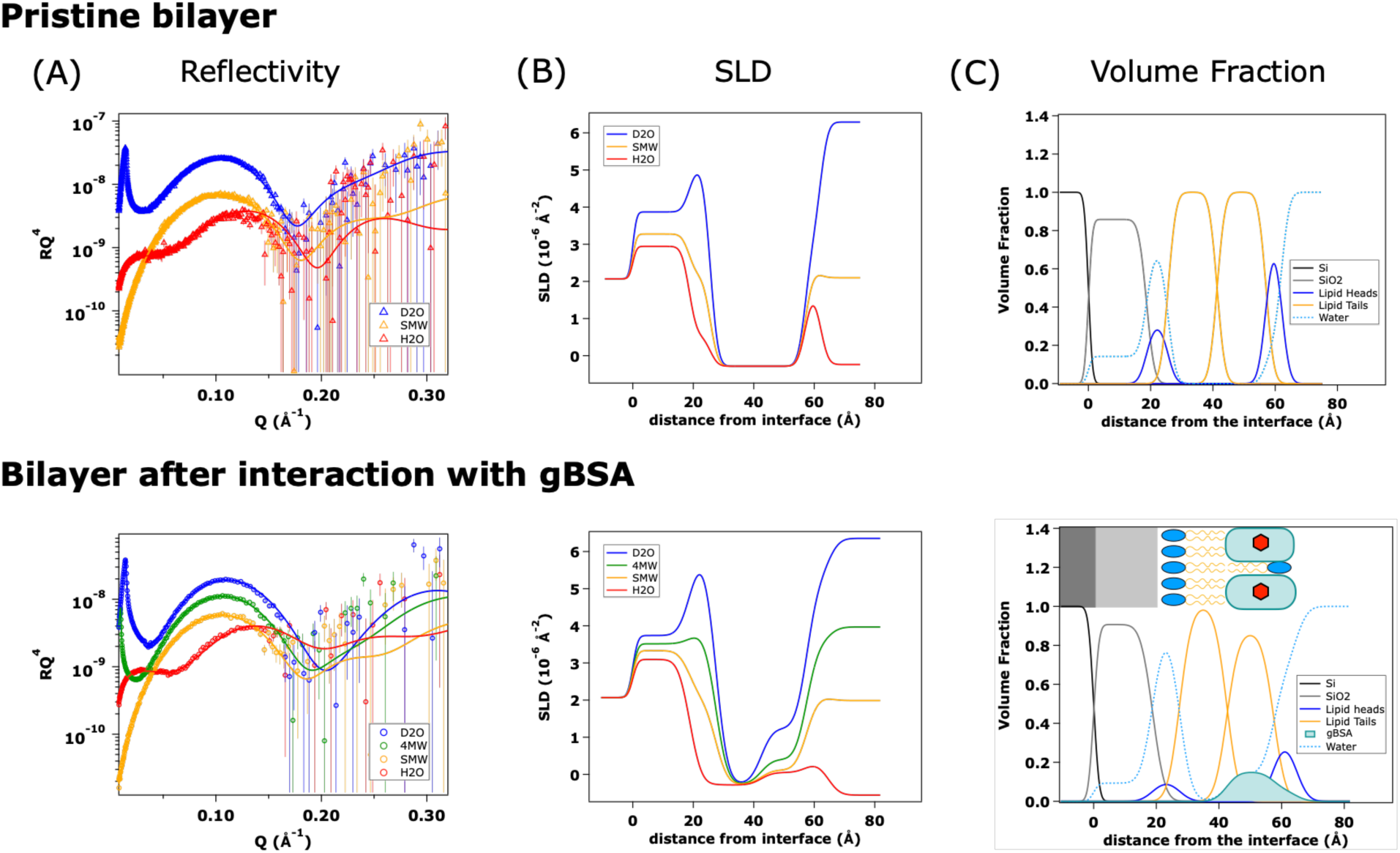
Reflectivity curves and fits at different contrasts (A), the SLDs obtained from the analysis (B) and the corresponding volume fraction distribution (C) of the SLB POPC/POPS 8:2 before and after the exposure to the gBSA. Symbols represent the raw data; continuous lines represent the fit. The model that describes best the experimental behaviour is the one representing the penetration of the protein in the outer leaflet of the lipid bilayer with no adsorbed layer of proteins onto the SLB.

**Table 2:**
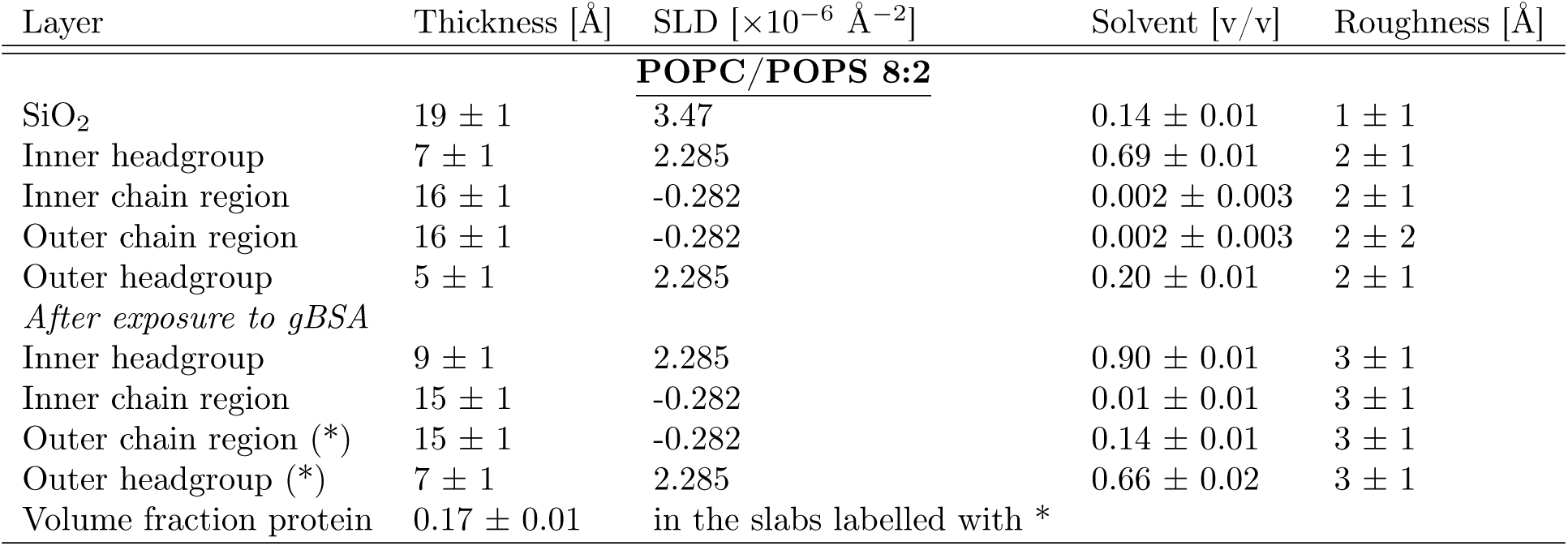
Structural parameters of the SLB POPC/POPS 8:2 before and after the exposure to the gBSA. The corresponding fits are in Figure 4. The values without error were kept fixed at their nominal value.

The membrane-associated volume fraction of gBSA is significantly higher than that of BSA, indicating enhanced interaction with negatively charged membranes. Although PS is predominantly localized to the inner leaflet, its externalization occurs in key processes such as platelet activation, coagulation, apoptosis, and in extracellular vesicles, ^46–49^ making its inclusion in our model membranes biologically relevant. The enhanced interaction of gBSA with PS-containing membrane may be attributed to glycation-induced changes in surface properties and charge distribution of the protein. Glycation primarily targets lysine and arginine residues, converting their positively charged side chains into neutral sugar adducts, and therefore reducing local positive charge. In theory, this modification would be expected to decrease electrostatic attraction to anionic lipids. However, the stronger interaction observed with gBSA may be explained by conformational rearrangements that alter the accessibility of hydrophilic or charged regions.

Glycation of BSA has been shown to cause destabilization or unfolding of the albumin structure, resulting in a 25–50% reduction in α-helix content and a 2-fold increase in β-sheets and turns structures.^50^ Molecular dynamics simulations further support these findings, showing that glycation of HSA induces conformational destabilization, increased flexibility, and altered hydration patterns that can affect protein dynamics and interactions. ^51^ Similarly, experimental studies on glycated BSA have demonstrated alterations in secondary structure and ligand-binding properties, such as reduced binding affinity toward gliclazide, highlighting the functional consequences of glycation-induced structural changes. ^52^ Such structural modifications can influence the protein’s interactions with other molecules, including lipid membranes, and may contribute to the differences observed between glycated and native albumin in our study.

Interestingly, previous studies have shown that glycation can increase the isoelectric point (pI) of BSA: for example, extensive glycation was found to increase the pI from 4.2 to 6.3. ^38^ Although this seems counterintuitive, such a shift may result from a combination of factors, including masking of acidic residues, glycation-induced changes in local electrostatic environments, and structural unfolding that exposes otherwise buried basic residues. Although the glycation yield in our study was substantially lower (five glucose moieties), even partial modification may be sufficient to affect the surface charge distribution and conformation of BSA in a way that enhances its interaction with negatively charged lipid membranes. Previous work has shown that glycation of serum albumin, particularly at lysine and arginine residues, alters protein conformation and ligand-binding characteristics.^53–56^ In another system, glycated *β*_2_-microglobulin exhibited altered binding to negatively charged dialysis membranes, suggesting that glycation may modulate protein–surface interactions through changes in molecular properties. ^57^

Moreover, glycation also introduces bulky, hydrophilic sugar groups that may strengthen interactions with lipid headgroups and the interfacial water layer. Previous work has shown that both glycosylation and glycation can modulate protein interactions, particularly in systems where surface chemistry and hydration dynamics are critical.^58–60^

## Conclusion

In this study, we performed a comparative analysis of bovine serum albumin (BSA) and its chemically-enhanced glycated form (gBSA) to investigate how glycation influences protein interactions with artificial lipid bilayers designed to mimic cellular membranes. Our results revealed that both BSA and gBSA interacted significantly with negatively charged SLB, with glycation enhancing this effect demonstrated by a substantial increase in membrane-associated protein volume fraction from 0.11 for BSA to 0.17 for gBSA. Neither BSA nor gBSA interacted significantly with zwitterionic and cationic lipid membranes.

These data provide molecular-level evidence that glycation strengthens albumin’s affinity for negatively charged membranes. While changes in surface charge distribution likely play a central role, increased hydrophilicity due to sugar additions may further enhance the binding. These findings could have implications for the biological behavior of glycated proteins in the bloodstream. If glycation enhances membrane or lipoprotein binding, it may affect the bioavailability, distribution, and ultimately, the diagnostic detectability of glycated albumin in serum assays. This could lead to underestimation or variability in clinical measurements, especially in hyperglycemic conditions where glycation levels are elevated.

The present study was designed as an initial step to underline the importance of glycation in these interactions. For future studies, it would be valuable to include phosphatidylinositol (PI), which represents approximately 10% of total phospholipids in mammalian plasma membranes,^61^ in future SLB models. PI is rich in hydroxyl groups which are capable of forming hydrogen bonds with proteins, an interaction avenue often undervalued compared ionic interactions. Incorporating PI into membrane models would better reflect physiological complexity, uncovering whether glycated albumin might form additional hydrogen bonds that reinforce the binding. These insights would enable more precise lipid-based biosensor designs by informing which lipid components should be avoided to minimize non-specific albumin adsorption. Additionally, a more precise control over glycation levels, both in terms of modification sites and glycation extent, would be valuable in discriminating the contribution of specific sugar moieties. Future experiments should also include higher protein concentrations to better mimic hyperglycemic conditions and extend findings to human serum albumin (HSA) for improved clinical relevance. Furthermore, future studies—including experiments with more tightly controlled levels of glycation, the use of complementary biophysical methods (dynamic light scattering and zeta potential performed on spherical vescicles), and molecular dynamics simulations could provide a more comprehensive mechanistic understanding of glycated albumin–membrane interactions.

Collectively, our findings deepen the understanding of how glycation modifies protein–lipid interactions and may guide the development of more accurate diagnostic and therapeutic strategies targeting glycation-related pathologies.

## Acknowledgement

We acknowledge the InnovaXN (European Union’s Horizon 2020 research and innovation programme under the Marie Skłodowska-Curie grant agreement no. 847439) for funding the InnovaXN-25-2020 project, the Institut Laue-Langevin that provided allocation of neutron beam time (Experiment DOI: doi:10.5291/ILL-DATA.9-13-1048, doi:10.5291/ILL-DATA.9-13-1084, doi:10.5291/ILL-DATA.9-13-1125) and the use of Partnership for Soft Condensed Matter laboratories. We also thank the Institut de Biologie Structurale (IBS) in Grenoble for performing the mass spectrometry analyses.

## Supporting Information Available

The following files are available free of charge: additional experimental data analysis including illustrative sketches of data modeling.

- SI.pdf: supporting information

