## Supplementary material for "Glycation enhances protein association with lipid bilayer membranes"

‖*ANSTO - Sydney, New Illawarra Rd, Lucas Heights NSW 2234, Australia*

⊥*Université Grenoble-Alpes, INSERM U13, CEA, Institute of Interdisciplinary Research of Grenoble (IRIG), Laboratory of Biosciences and Bioengineering for Health (BGE)-BIOMICS, 38054 Grenoble, France*

#*Université Grenoble-Alpes, CNRS, Grenoble INP, LMGP-UMR 5628, 38000 Grenoble, France*

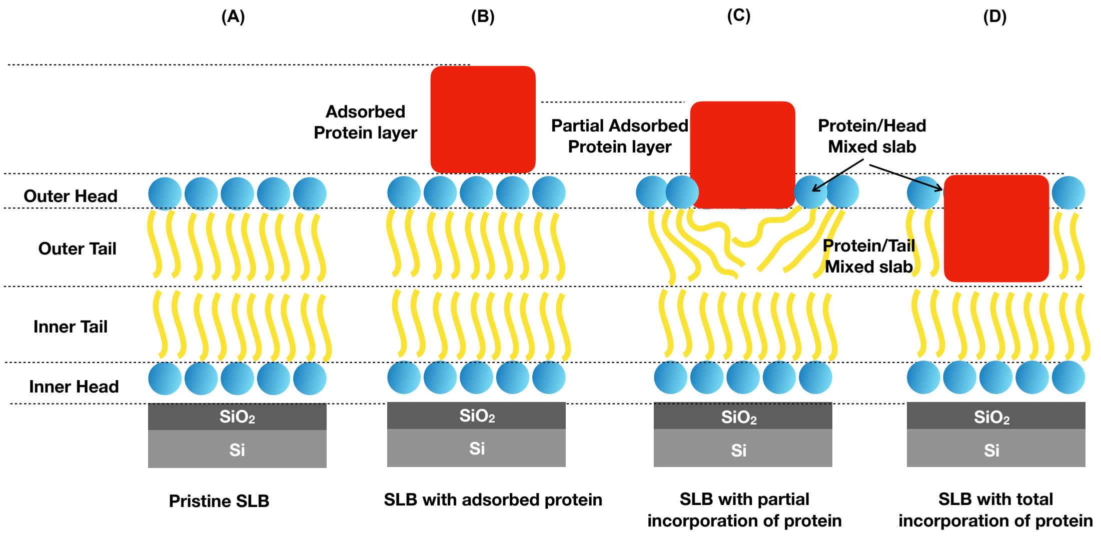


Figure S1: *Slab models used to build the model for the analysis of the reflectivity data.*

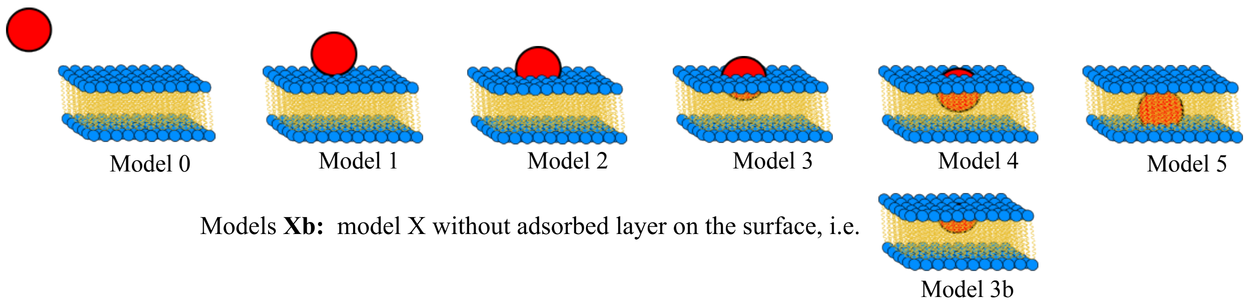


Figure S2: *Schematic representations of all the models tested describing the possible interactions between a protein and a SLB. The red circle represents schematically the average position of the protein with respect to the SLB. model 0 describes lacks of protein on the lipid bilayer; model 1 describes adsorption of the protein onto the SLB; model 2 describes adsorption of the protein onto the SLB and penetration into the head group layer of the outer lipid leaflet; model 3 describes adsorption of the protein onto the SLB and penetration into the outer leaflet; model 4 describes adsorption of the protein onto the SLB and penetration into the outer leaflet and inner tail of the membrane; and model 5 describes the total incorporation of the protein in the SLB. Additionally, we tested models defined as Xb, where X could be any model between 2 and 5, corresponding to each model without the presence of the protruding adsorbed protein layer on the surface of the SLB (*S2*, model 3b).*

Table S1: Normalized $\chi^2$ values for the different models tested for pure POPC lipid bilayer after exposure to BSA and gBSA. The chosen model is highlighted in the table. It is important to note that the best model was chosen accordingly to the smallest normalized $\chi^2$ value but also with a further discrimination using the Bayesian analysis. However, due to the relatively small magnitude of these differences, an unambiguous model selection was not achievable.

| Protein | Model 0 | Model 1 | Model 2 | Model 3 | Model 4 | Model 5 |
|---|---|---|---|---|---|---|
| BSA | 3.01 | 2.91 | 2.99 | 3.64 | 4.79 | 5.06 |
| gBSA | 3.98 | 3.62 | 3.61 | 5.02 | 6.31 | 7.15 |

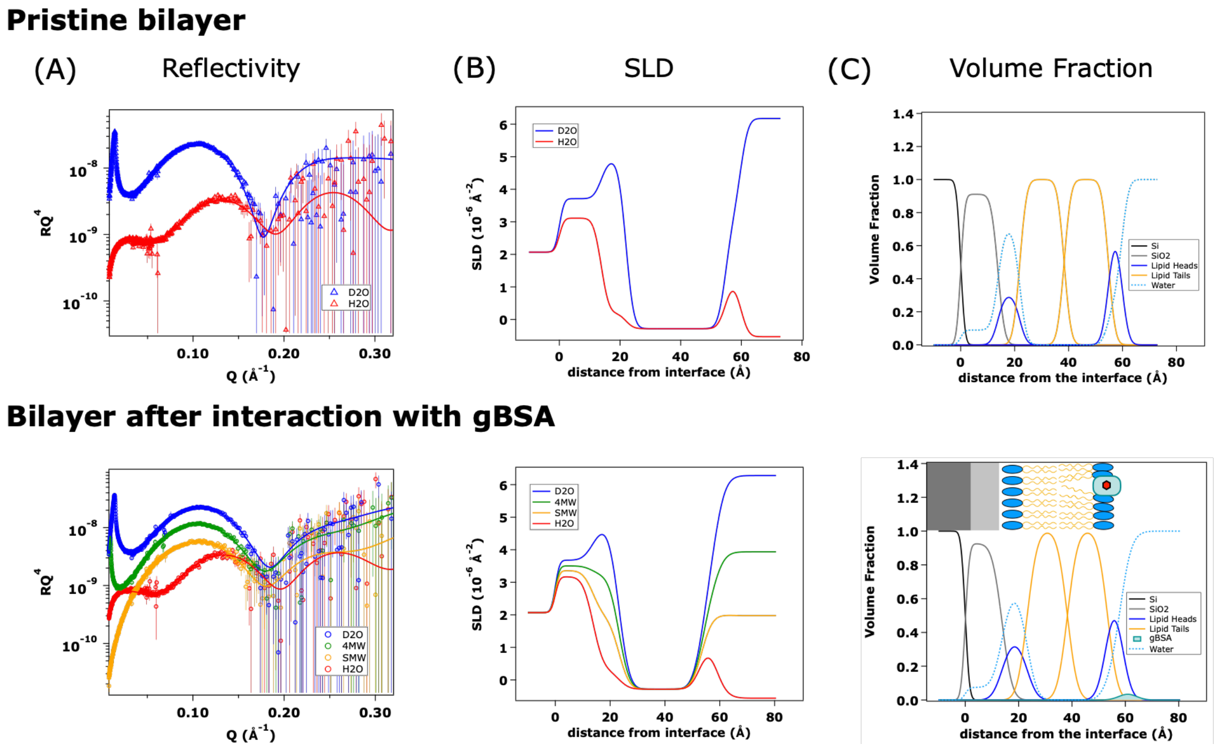


Figure S3: *Reflectivity curves and fits at different contrasts (A), the SLDs obtained from the analysis (B) and the corresponding volume fraction distribution (C) of the SLB POPC before and after the exposure to the gBSA. Symbols represent the raw data; continuous lines represent the fits obtained from the model. The represented model is the one showing protein penetration into the outer head group region of the lipid bilayer with an additional protruding protein layer above the bilayer.*

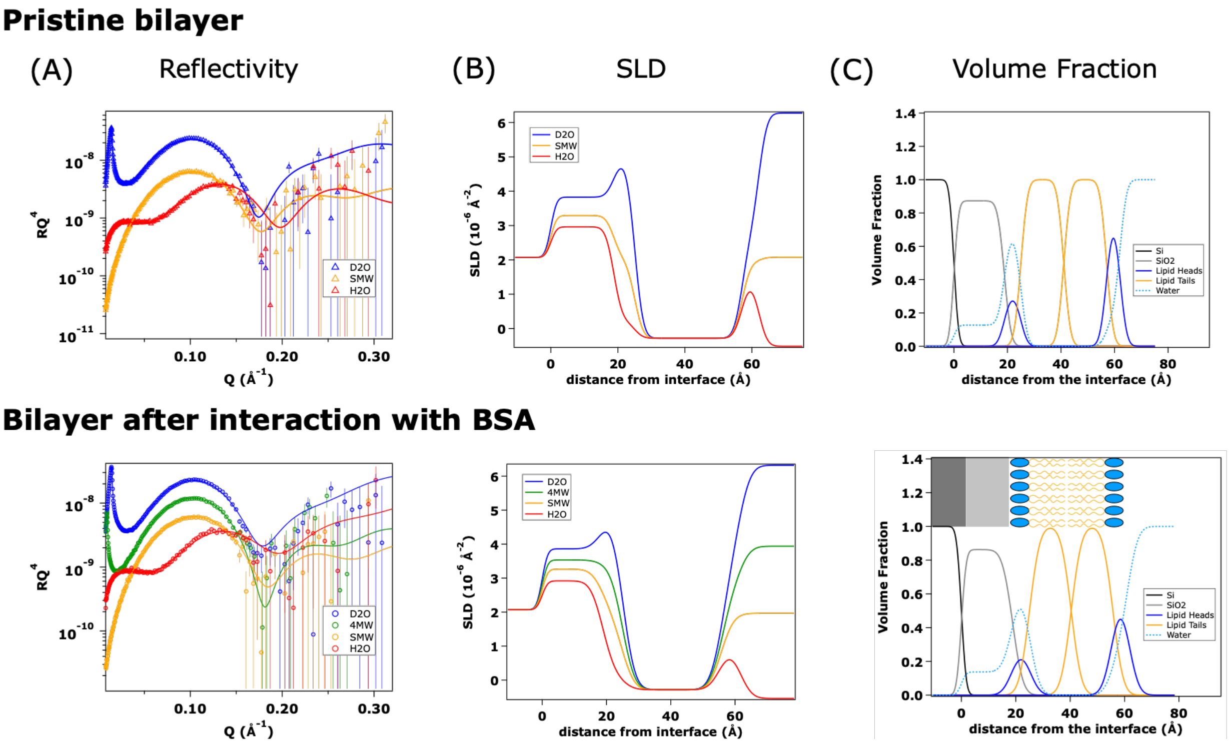


Figure S4: *Reflectivity curves and fits at different contrasts (A), the SLDs obtained from the analysis (B) and the corresponding volume fraction distribution (C) of the SLB POPC before and after the exposure to the BSA. Symbols represent the raw data; continuous lines represent the fit. The model that describes better the experimental behaviour is the model representing no interaction between the protein and the lipid bilayer.*

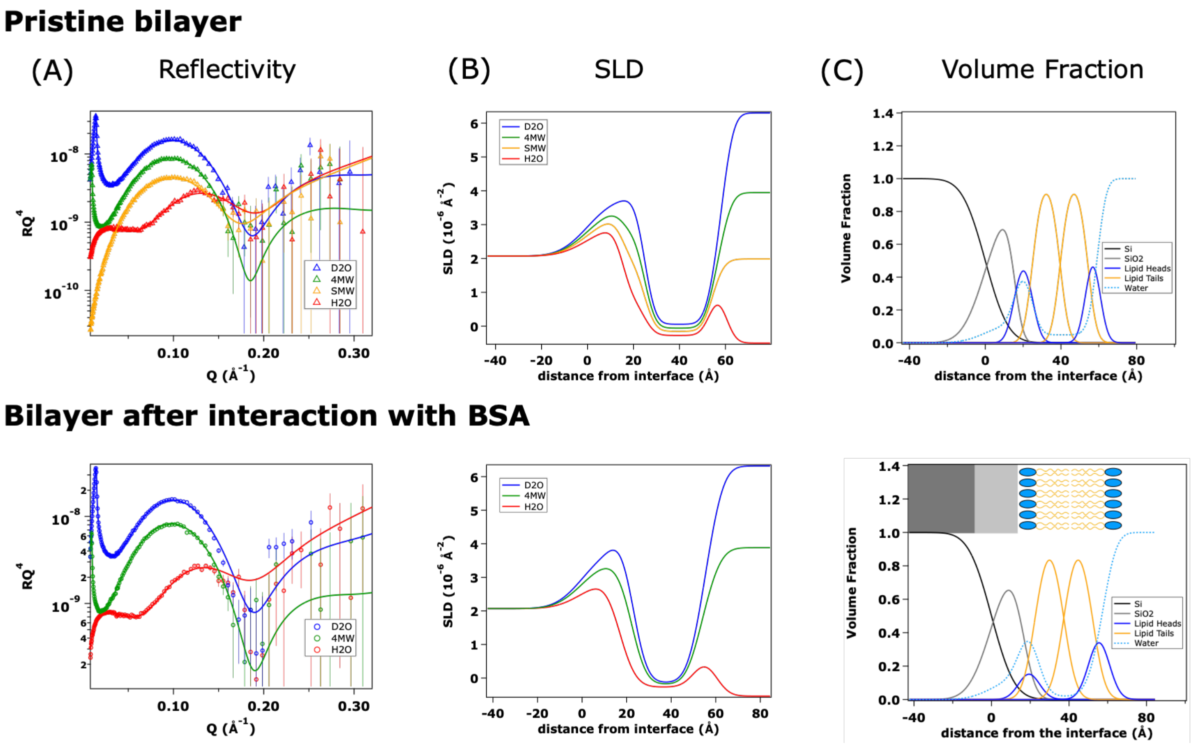


Figure S5: *Reflectivity curves and fits at different contrasts (A), the SLDs obtained from the analysis (B) and the corresponding volume fraction distribution (C) of the SLB POPC/DOTAP 7:3 before and after the exposure to the BSA. Symbols represent the raw data; continuous lines represent the fit. The model that describes better the experimental behaviour is the model representing no interaction between the protein and the lipid bilayer.*

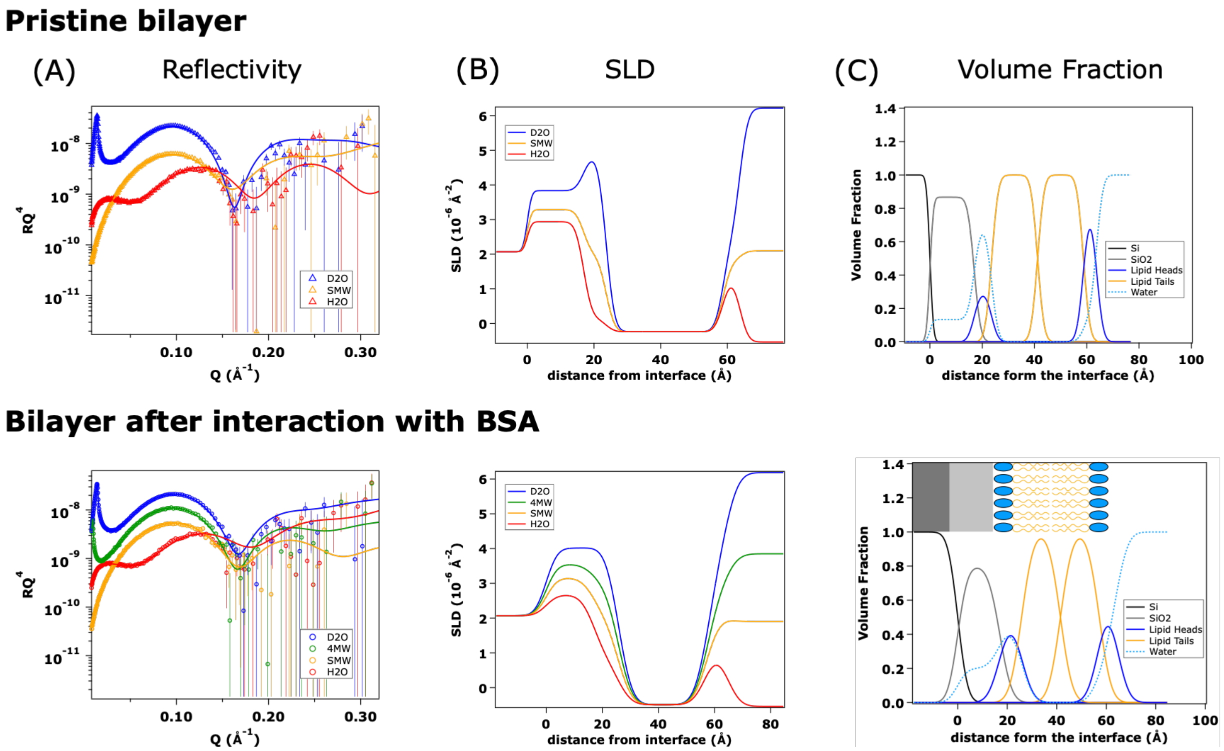


Figure S6: *Reflectivity curves and fits at different contrasts (A), the SLDs obtained from the analysis (B) and the corresponding volume fraction distribution (C) of the SLB POPC/SM/CHOL 6:3:1 before and after the exposure to the BSA. Symbols represent the raw data; continuous lines represent the fit. The model that describes better the experimental behaviour is the model representing no interaction between the protein and the lipid bilayer.*

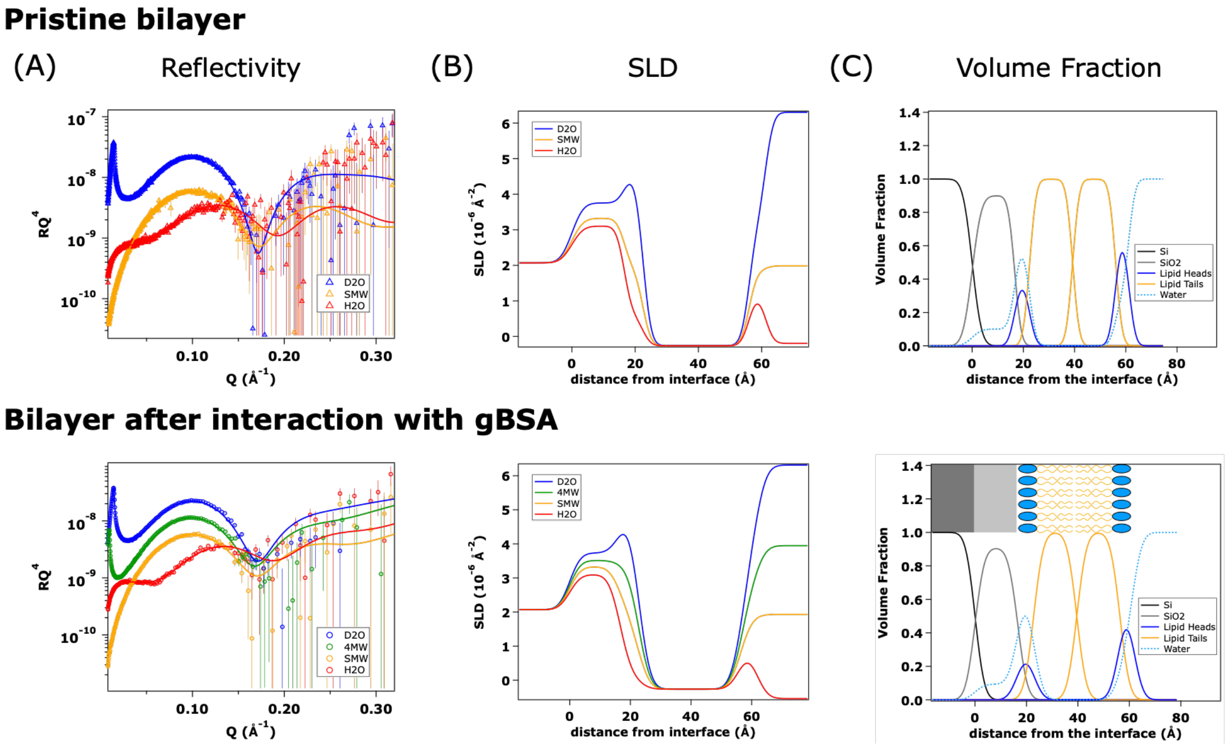


Figure S7: *Reflectivity curves and fits at different contrasts (A), the SLDs obtained from the analysis (B) and the corresponding volume fraction distribution (C) of the SLB POPC/DOTAP 7:3 before and after the exposure to the gBSA. Symbols represent the raw data; continuous lines represent the fit. The model that describes better the experimental behaviour is the one representing no interaction between the protein and the lipid bilayer.*

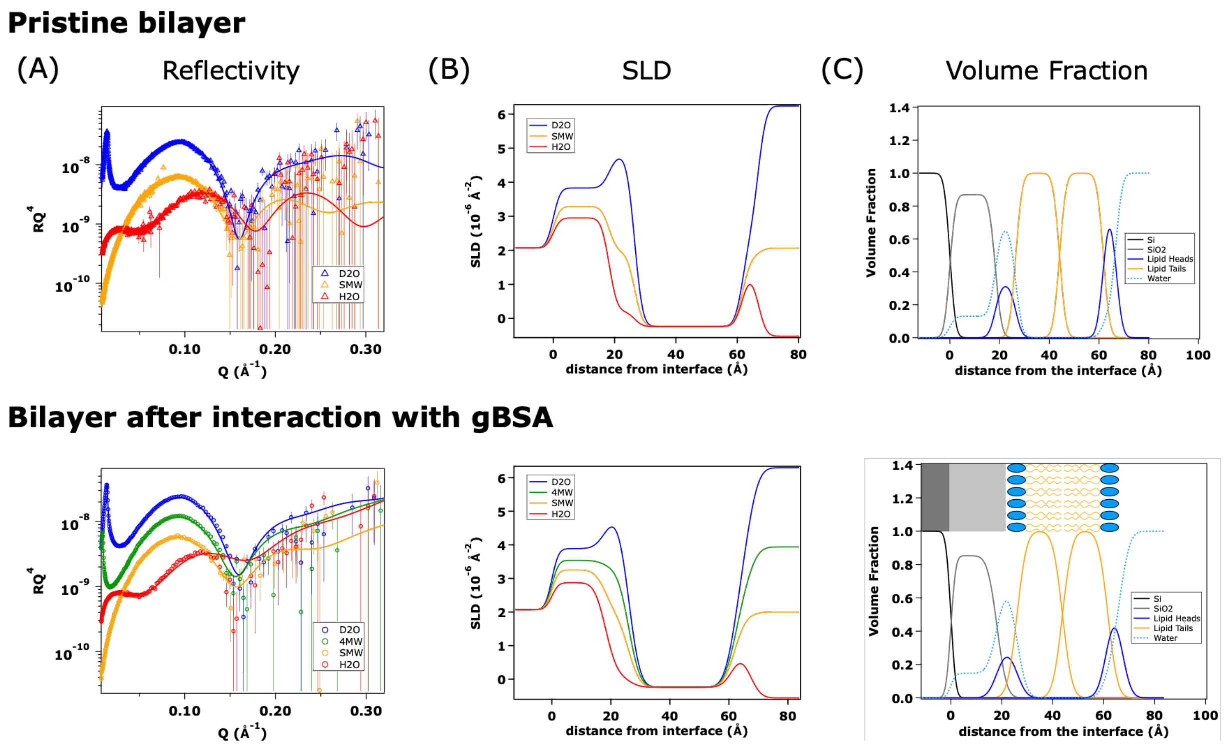


Figure S8: *Reflectivity curves and fits at different contrasts (A), the SLDs obtained from the analysis (B) and the corresponding volume fraction distribution (C) of the SLB POPC/SM/CHOL 6:3:1 before and after the exposure to the gBSA. Symbols represent the raw data; continuous lines represent the fit. The model that describes better the experimental behaviour is the one representing no interaction between the protein and the lipid bilayer.*

Table S2: *Structural parameters of the SLBs before and after the exposure to the gBSA. The corresponding fits are in Figures* S*3,* S*7 and* S*8. The values without error were kept fixed at their nominal value.*

| Layer | Thickness [Å] | SLD [$\times10^{-6}$ Å$^{-2}$] | Solvent [v/v] | Roughness [Å] |
|---|---|---|---|---|
| | | **POPC** | | |
| $SiO_2$ | 14 ± 1 | 3.47 | 0.09 ± 0.01 | 1 ± 1 |
| Inner headgroup | 8 ± 1 | 1.88 | 0.70 ± 0.01 | 2 ± 1 |
| Inner chain region | 16 ± 1 | -0.28 | 0.0001 ± 0.0003 | 2 ± 1 |
| Outer chain region | 16 ± 1 | -0.28 | 0.0001 ± 0.0003 | 2 ± 1 |
| Outer headgroup | 5 ± 1 | 1.88 | 0.28 ± 0.01 | 2 ± 1 |
| *After exposure to gBSA* | | | | |
| Inner headgroup | 9 ± 1 | 1.88 | 0.64 ± 0.01 | 3 ± 1 |
| Inner chain region | 15 ± 1 | -0.28 | 0.00 ± 0.01 | 3 ± 1 |
| Outer chain region | 15 ± 1 | -0.28 | 0.00 ± 0.01 | 3 ± 1 |
| Outer headgroup (*) | 5 ± 1 | 1.88 | 0.21 ± 0.01 | 3 ± 1 |
| Protein layer (*) | 5 ± 1 | | 0.94 ± 0.01 | 3 ± 1 |
| Volume fraction protein | 0.06 ± 0.01 | in the slabs labelled with * | | |
| | | **POPC/DOTAP 7:3** | | |
| $SiO_2$ | 17 ± 1 | 3.47 | 0.10 ± 0.01 | 3 ± 1 |
| Inner headgroup | 6 ± 1 | 1.796 | 0.61 ± 0.01 | 2 ± 1 |
| Inner chain region | 17 ± 1 | -0.262 | 0.001 ± 0.001 | 2 ± 1 |
| Outer chain region | 17 ± 1 | -0.262 | 0.001 ± 0.001 | 2 ± 1 |
| Outer headgroup | 5 ± 1 | 1.796 | 0.29 ± 0.01 | 2 ± 1 |
| *After exposure to gBSA* | | | | |
| Inner headgroup | 6 ± 1 | 1.796 | 0.70 ± 0.01 | 3 ± 1 |
| Inner chain region | 17 ± 1 | -0.262 | 0.001 ± 0.001 | 3 ± 1 |
| Outer chain region | 17 ± 1 | -0.262 | 0.001 ± 0.001 | 3 ± 1 |
| Outer headgroup | 5 ± 1 | 1.796 | 0.30 ± 0.01 | 3 ± 1 |
| | | **POPC/SM/CHOL 6:3:1** | | |
| $SiO_2$ | 18 ± 1 | 3.47 | 0.13 ± 0.01 | 2 ± 1 |
| Inner headgroup | 8 ± 1 | 1.725 | 0.67 ± 0.01 | 2 ± 1 |
| Inner chain region | 18 ± 1 | -0.238 | 0.0007 ± 0.0001 | 2 ± 1 |
| Outer chain region | 18 ± 1 | -0.238 | 0.0007 ± 0.0001 | 2 ± 1 |
| Outer headgroup | 5 ± 1 | 1.725 | 0.15 ± 0.003 | 2 ± 1 |
| *After exposure to gBSA* | | | | |
| Inner headgroup | 8 ± 1 | 1.725 | 0.70 ± 0.01 | 3 ± 1 |
| Inner chain region | 18 ± 1 | -0.238 | 0.0001 ± 0.0001 | 3 ± 1 |
| Outer chain region | 18 ± 1 | -0.238 | 0.0001 ± 0.0001 | 3 ± 1 |
| Outer headgroup | 5 ± 1 | 1.725 | 0.30 ± 0.01 | 3 ± 1 |

Table S3: *Structural parameters of the SLBs before and after the exposure to BSA. The corresponding fits are in Figures S4, S5 and S6. The values without error were kept fixed at their nominal value.*

| Layer | Thickness [Å] | SLD [$\times 10^{-6}$ Å$^{-2}$] | Solvent [v/v] | Roughness [Å] |
|---|---|---|---|---|
| | | **POPC** | | |
| $SiO_2$ | 19 ± 1 | 3.47 | 0.13 ± 0.01 | 1 ± 1 |
| Inner headgroup | 6 ± 1 | 1.88 | 0.69 ± 0.01 | 2 ± 1 |
| Inner chain region | 16 ± 1 | -0.28 | 0.0001 ± 0.0001 | 2 ± 1 |
| Outer chain region | 16 ± 1 | -0.28 | 0.0001 ± 0.0001 | 2 ± 1 |
| Outer headgroup | 5 ± 1 | 1.88 | 0.18 ± 0.01 | 2 ± 1 |
| *After exposure to BSA* | | | | |
| Inner headgroup | 6 ± 1 | 1.88 | 0.69 ± 0.01 | 3 ± 1 |
| Inner chain region | 15 ± 1 | -0.28 | 0.004 ± 0.001 | 3 ± 1 |
| Outer chain region | 15 ± 1 | -0.28 | 0.004 ± 0.001 | 3 ± 1 |
| Outer headgroup | 6 ± 1 | 1.88 | 0.30 ± 0.01 | 3 ± 1 |
| | | **POPC/DOTAP 7:3** | | |
| $SiO_2$ | 16 ± 1 | 3.47 | 0.11 ± 0.01 | 9 ± 1 |
| Inner headgroup | 8 ± 1 | 1.796 | 0.52 ± 0.01 | 4 ± 1 |
| Inner chain region | 15 ± 1 | -0.262 | 0.03 ± 0.01 | 4 ± 1 |
| Outer chain region | 15 ± 1 | -0.262 | 0.03 ± 0.01 | 4 ± 1 |
| Outer headgroup | 5 ± 1 | 1.796 | 0.11 ± 0.01 | 4 ± 1 |
| *After exposure to BSA* | | | | |
| Inner headgroup | 6 ± 1 | 1.796 | 0.66 ± 0.01 | 5 ± 1 |
| Inner chain region | 15 ± 1 | -0.262 | 0.02 ± 0.01 | 5 ± 1 |
| Outer chain region | 15 ± 1 | -0.262 | 0.02 ± 0.01 | 5 ± 1 |
| Outer headgroup | 7 ± 1 | 1.796 | 0.30 ± 0.01 | 5 ± 1 |
| | | **POPC/SM/CHOL 6:3:1** | | |
| $SiO_2$ | 17 ± 1 | 3.47 | 0.13 ± 0.01 | 1 ± 1 |
| Inner headgroup | 7 ± 1 | 1.725 | 0.70 ± 0.01 | 2 ± 1 |
| Inner chain region | 18 ± 1 | -0.238 | 0.0007 ± 0.0001 | 2 ± 1 |
| Outer chain region | 18 ± 1 | -0.238 | 0.0007 ± 0.0001 | 2 ± 1 |
| Outer headgroup | 5 ± 1 | 1.725 | 0.15 ± 0.003 | 2 ± 1 |
| *After exposure to BSA* | | | | |
| Inner headgroup | 7 ± 1 | 1.753 | 0.70 ± 0.01 | 3 ± 1 |
| Inner chain region | 17 ± 1 | -0.238 | 0.0001 ± 0.0001 | 3 ± 1 |
| Outer chain region | 17 ± 1 | -0.238 | 0.0001 ± 0.0001 | 3 ± 1 |
| Outer headgroup | 6 ± 1 | 1.753 | 0.30 ± 0.01 | 3 ± 1 |

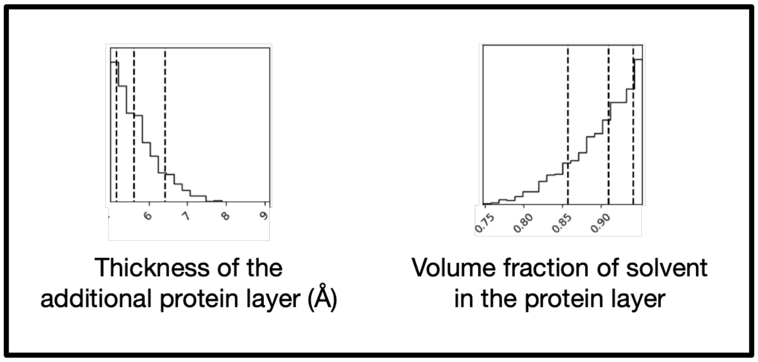


Figure S9: *Posterior distribution of the key parameters of the additional protein layer obtained from the Bayesian analysis of BSA interacting with the POPC/POPS supported bilayer. The key parameters are the volume fraction of solvent and the thickness of the protein layer.*
